# Post-stroke administration of the p75 neurotrophin receptor modulator, LM11A-31, attenuates chronic changes in brain metabolism, increases neurotransmitter levels, and improves recovery

**DOI:** 10.1101/2021.04.30.442181

**Authors:** Thuy-Vi V. Nguyen, Rachel H. Crumpacker, Kylie E. Calderon, Frankie G. Garcia, Jacob C. Zbesko, Jennifer B. Frye, Selena Gonzalez, Danielle A. Becktel, Tao Yang, Marco A. Tavera-Garcia, Helena W. Morrison, Rick G. Schnellmann, Frank M. Longo, Kristian P. Doyle

## Abstract

The aim of this study was to test whether post-stroke oral administration of a small molecule p75 neurotrophin receptor (p75^NTR^) modulator (LM11A-31) can augment neuronal survival and improve recovery in a mouse model of stroke. Mice were administered LM11A-31 for up to 12 weeks, beginning 1 week after stroke. Metabolomic analysis revealed that after 2 weeks of daily treatment, mice that received LM11A-31 were distinct from vehicle treated mice by principal component analysis and had higher levels of serotonin, acetylcholine, and dopamine in their ipsilateral hemisphere. LM11A-31 treatment also improved redox homeostasis by restoring reduced glutathione. It also offset a stroke induced reduction in glycolysis by increasing acetyl-CoA. There was no effect on cytokine levels in the infarct. At 13 weeks following stroke, adaptive immune cell infiltration in the infarct was unchanged in LM11A-31 treated mice, indicating that LM11A-31 does not alter the chronic inflammatory response to stroke at the site of the infarct. However, LM11A-31 treated mice had less brain atrophy, neurodegeneration, tau pathology, and microglial activation in other regions of the ipsilateral hemisphere. These findings correlated with improved recovery of motor function on a ladder test, improved sensorimotor and cognitive abilities on a nest construction test, and less impulsivity in an open field test. These data support small molecule modulation of the p75 neurotrophin receptor for preserving neuronal health and function during stroke recovery.

**SIGNIFICANCE STATEMENT:** The findings from this study introduce the p75 neurotrophin receptor as a novel small molecule target for promotion of stroke recovery. Given that LM11A-31 is in clinical trials as a potential therapy for Alzheimer’s disease, it could be considered as a candidate for assessment in stroke or vascular dementia studies.

## INTRODUCTION

People over the age of 55 have a 1 in 6 lifetime risk of having a stroke (Seshadri et al., 2006) and there are more than 6.5 million survivors of stroke in the United States (US) (http://www.strokecenter.org/patients/about-stroke/stroke-statistics/; Donkor, 2018). The only FDA approved drug available for the treatment of stroke is tissue plasminogen activator (TPA), but it must be administered within 4.5 hours of ischemic stroke onset. Mechanical thrombectomy can also be used to restore cerebral blood flow in acute ischemic stroke, however, less than 20% of patients qualify for this type of reperfusion therapy and it must be initiated within 24 hours of stroke onset (Rabinstein, 2020). In contrast, no pharmacologic therapies are available to enhance recovery in the days and weeks after stroke. This prompts the need to identify compounds commonly applied to, or in the pipeline for, the treatment of other neurological disorders to determine if they can promote neural repair and recovery of cellular function after stroke. One such example is LM11A-31, a small molecule ligand for the p75 neurotrophin receptor (p75^NTR^). LM11A-31 is currently under Phase IIa clinical evaluation for the treatment of mild to moderate Alzheimer’s disease (AD; ClinicalTrials.gov Identifier: NCT03069014).

p75^NTR^ is expressed in basal forebrain cholinergic, cortical, hippocampal, noradrenergic, and other central nervous system (CNS) neurons (Becker et al., 2018) that have critical roles in learning and memory, cognition, psychological wellbeing, and sensorimotor functions. It is a tumor necrosis factor (TNF) family member receptor that recognizes neurotrophins, including nerve growth factor, neurotrophin-3, neurotrophin-4/5, and brain derived neurotrophic factor (BDNF). Although once categorized as a death initiating receptor, it is now clear that it can also promote cell survival through phosphoinositide 3-kinase (PI3K)/protein kinase B (AKT) and interleukin-1 receptor-associated kinase (IRAK)/nuclear factor kappa B (NFκB) dependent signaling mechanisms (Longo and Massa, 2008). The balance of pro-survival versus pro-apoptotic signaling is determined by the availability of adaptor molecules, which is itself determined by cell type and contextual cues (Longo and Massa, 2008).

LM11A-31 specifically functions as a p75^NTR^ ligand. This specificity is supported by multiple lines of evidence, including the rapid recruitment of the IRAK p75^NTR^ intracellular adaptor following treatment *in vitro*, the ability of LM11A-31 to block a p75^NTR^ antibody from binding to the receptor, and conversely the ability of p75^NTR^ antibodies to block LM11A-31 activity. It is also supported by loss of ligand signaling and function in p75^NTR-/-^ neurons, nerve growth factor (NGF) like promotion of p75^NTR^ endocytosis, and no detectable effect of LM11A-31 on tropomyosin receptor A (TrkA) (Longo and Massa, 2008; Longo and Massa, 2013). At nanomolar concentrations *in vitro*, LM11A-31 prevents neuronal death, promotes neurotrophic signaling, and inhibits neurodegenerative signaling through the p75^NTR^ (Massa et al., 2006; Yang et al., 2008). Therefore, LM11A-31 functions as a p75^NTR^ receptor modulator rather than a simple agonist or antagonist, switching p75^NTR^ from neurodegenerative to neurotrophic signaling in neurons expressing p75^NTR^ (Xie et al., 2019).

Importantly, LM11A-31 has shown therapeutic effects in multiple animal models of neurological diseases, including post-traumatic brain injury (Shi et al., 2013), human immunodeficiency virus (HIV) related dementia (Meeker et al., 2016; Fogle et al., 2021; Xie et al., 2021), contusion spinal cord injury (Tep et al., 2013), Huntington’s disease (Simmons et al., 2016; Simmons et al., 2021), and AD (Knowles et al., 2013; Nguyen et al., 2014; Simmons et al., 2014; Yang et al., 2020a). In AD, LM11A-31 has been shown to be efficacious in male and female mice and in two different mouse models of AD (Simmons et al., 2014). It has also been shown to reverse neuronal degeneration in late stage AD pathology in transgenic mice (Knowles et al., 2009; Knowles et al., 2013; Simmons et al., 2013; Nguyen et al., 2014; James et al., 2017). LM11A-31 is orally bioavailable and CNS penetrant, exerting maximal efficacy in the 25-50 mg/kg range (HCl salt form), which at 50 mg/kg corresponds to a human equivalent dose of 284 mg for a 70 kg adult, per body surface area mouse to human conversion (Massa et al., 2006; Yang et al., 2008). LM11A-31 reaches and preferentially accumulates in the brain compared to blood (Knowles et al., 2013), which reduces the likelihood of systemic toxicity.

In light of the efficacy of LM11A-31 across a wide spectrum of neurological diseases, its attractive pharmacokinetic properties, and the fact that it is currently undergoing clinical testing for the treatment of AD, we sought to determine whether LM11A-31 could protect neurons from the chronic sequelae of stroke that cause neurodegeneration in the weeks after stroke. We discovered in a mouse model of experimental stroke that daily oral administration of LM11A-31 can mitigate secondary neurodegeneration, increase neurotransmitter levels, and improve functional recovery. These findings correlated with changes in redox homeostasis and energy metabolism in the injured hemisphere, but no changes in inflammation within the infarct. Together, these data support the use of LM11A-31 for preserving neuronal health and function in patients recovering from stroke.

## MATERIALS AND METHODS

### Mice

17-18-month-old, male C57BL/6J (The Jackson Laboratory, Stock No: 000664) mice were used for all experiments. All mice were given *ad libitum* access to food and water and were maintained in a 12-hour light/dark cycle in a temperature controlled housing facility. In accordance with Stroke Therapy Academic Industry Roundtable (STAIR) recommendations (Stroke Therapy Academic Industry, 1999), mice from Phase I (Cohort 1) were randomly allocated into treatment groups before the stroke/sham procedure. Additionally, mice from Phase II (Cohort 2) were stroked and stratified into treatment groups based on Day 2 post-stroke ladder rung data. All dosing, behavioral experiments, imaging, and morphological and biochemical analyses were performed with blinding relative to vehicle versus LM11A-31 treatment. All procedures met National Institute of Health guidelines with the approval of the University of Arizona Institutional Animal Care and Use Committee.

### Stroke and sham surgeries

Distal Middle Cerebral Artery Occlusion (dMCAO) with hypoxia (DH stroke) was performed as previously published (Doyle et al., 2012). Briefly, following anesthesia by isoflurane inhalation, the temporalis muscle was exposed by incision. The right MCA was identified and exposed using a microdrill on the skull. The meninges were cut, and the MCA was cauterized. The incision was then closed using surgical glue. Directly after receiving the dMCAO, mice underwent hypoxia (9% oxygen and 91% nitrogen) for 45 minutes. Sham operated mice underwent the same surgical steps, except for cauterization of the dMCA. They were also given 9% oxygen and 91% nitrogen for 45 minutes immediately after surgery. Throughout the surgery and hypoxia, mouse core temperature was monitored and maintained at 37°C. The purpose of hypoxia in this model is to increase infarct size, as C57BL/6 mice that receive a dMCAO without hypoxia have significantly smaller infarcts (Doyle et al., 2012). This stroke model results in a consistent infarct volume of approximately 25% of the ipsilateral hemisphere located in the somatosensory cortex and the M1 region of the motor cortex (Doyle et al., 2012; Nguyen et al., 2018). Mice were euthanized at 3 and 13 weeks following surgery with isoflurane inhalation and exsanguination and were perfused intracardially with 0.9% saline without heparin.

### LM11A-31

LM11A-31 [2-amino-3-methyl-pentanoic acid (2-morpholin-4-yl-ethyl)-amide], an isoleucine derivative, was synthesized by Ricerca Biosciences at >97% purity, as assessed by liquid chromatography/mass spectroscopy. LM11A-31 (−31; final dose: 50 mg/kg, HCl salt form) was prepared in sterile water and administered by oral gavage. This dose is efficacious in both male and female mice, in two models of AD, and has high efficacy on mid- to late-stage AD progression (Knowles et al., 2013; Nguyen et al., 2014; Simmons et al., 2014). An equivalent volume (10 μl per gram of body weight) of sterile water was administered by oral gavage to vehicle (veh) treated mice.

### Global metabolic profiling

At 3 weeks following surgery, infarcts from stroked mice, and the equivalent region of the cortex from sham operated mice, were resected and the remainder of the ipsilateral hemispheres were snap frozen and sent to Metabolon (Morrisville, NC) for processing and analysis (Ribbenstedt et al., 2018). At Metabolon, samples were prepared by first removing proteins and extracting small molecules and metabolites through a series of aqueous and organic extractions. The resulting extract was divided into 4 aliquots, each of which then underwent metabolic profiling. Two aliquots were profiled using reverse phase ultrahigh performance liquid chromatography/tandem mass spectrometry (UPLC-MS/MS) in acidic positive ion conditions, 1 aliquot in conditions optimized for more hydrophobic compounds and the other in conditions optimized for more hydrophilic compounds. Another aliquot was profiled using UPLC-MS/MS in basic negative ion conditions, and the 4^th^ aliquot using hydrophilic interaction ultra-performance liquid chromatography (HILIC/ULPC-MS/MS). Data extraction, compound identification, metabolite quantification, and data normalization were performed using Metabolon’s proprietary software. Quality control and curation procedures were used to remove artifacts, mis-assignments, and background noise (Choi et al., 2018).

### Multiplex immunoassay

At 3 weeks following surgery, infarcts from stroked mice and the equivalent region of the cortex from sham operated mice were dissected and sonicated in ice cold 0.1 M phosphate buffered saline containing 1% triton X-100 and 0.1% sodium deoxycholate, Protease Inhibitor Cocktail (1:100; Millipore Sigma), and Phosphatase Inhibitor Cocktail 2 (1:100; Millipore Sigma). Following centrifugation, the total protein concentration of each supernatant was measured using a Direct Detect Infrared Spectrometer (Millipore Sigma). Cytokines/chemokines were then quantified by multiplex immunoassay using a high sensitivity mouse multiplex magnetic bead kit (Millipore Sigma) and performed according to the manufacturer’s recommendations (MILLIPLEX MAP Mouse Cytokine/Chemokine Magnetic Bead Panel-Immunology Multiplex Assay, MCYTOMAG-70K). Each lysate sample, standard, and quality control were measured in duplicate. Plates were read using a MAGPIX instrument (Luminex, Austin, TX), and results were analyzed using MILLIPLEX Analyst 5.1 software (Millipore Sigma). For those analytes that were below the lower limit of detection, the lower limit of detection values was used for data analysis. Concentrations of cytokines/chemokines were normalized to concentrations of total protein in the lysates.

### Processing brain tissue for histology and immunohistochemistry

At 13 weeks following stroke, brains were post-fixed in 4% paraformaldehyde for 24 hours, followed by cryopreservation in 30% sucrose. Frozen coronal sections (40 μm) were then taken through the entire brain using a freezing Microtome HM 450 sliding microtome (ThermoFisher Scientific, Waltham, MA). Brain sections were sequentially placed into 16 microcentrifuge tubes and stored in cryoprotectant medium at –20°C until staining.

### Nissl staining

For Nissl staining, sections were mounted, dehydrated and cleared in xylenes, rehydrated, immersed in 0.5% cresyl violet acetate (MilliporeSigma, Burlington, MA) solution for 1 minute, rinsed with distilled water, differentiated in 0.25% acetic alcohol, dehydrated though a graded ethanol series, cleared, and coverslipped with mounting media.

### Immunohistochemistry

For immunostaining, sections were immunostained according to standard protocols (Nguyen et al., 2014). The following antibodies were used: mouse anti-neuronal nuclei (NeuN; Millipore Sigma), goat anti-choline acetyl transferase (ChAT; Millipore Sigma), rabbit anti-tyrosine hydroxylase (TH; Millipore Sigma), mouse anti-AT8 [recognizes tau phosphorylated at serine 202 and threonine 205 (p-tau^Ser202/Thr205^; ThermoFisher Scientific)], rat anti-CD68 (Bio-Rad AbD Serotec, Hercules, CA), rat anti-mouse B220/CD45R biotinylated (BD Biosciences, San Jose, CA), and hamster anti-CD3e (BD Biosciences). Briefly, free floating sections were immunolabeled with antibodies (1:500 for NeuN, 1:600 for ChAT, 1:1000 for TH, 1:800 for AT8, 1:1000 for CD68, 1:500 for B220/CD45R, and 1:1000 for CD3e) in conjunction with Mouse on Mouse (M.O.M.) Detection kit (for AT8), ABC Vector Elite, and 3,3’-diaminobenzidine (Vector Laboratories, Burlingame, CA) for visualization.

### Microscopy and imaging

Microscopy was performed by an experimenter blinded to treatment group. Captured images were analyzed by an independent experimenter also blinded to treatment group. Images shown in the figures are representative images that best depict the data represented by the graphs. Images were captured with a Keyence (Elmwood Park, NJ) BZ-X700 digital microscope or a Leica (Buffalo Grove, IL) DM6000B light microscope coupled with a Leica DFC 7000 T camera. The Mouse Brain in Stereotaxic Coordinates of Franklin and Paxinos 3^rd^ edition (Franklin and Paxinos, 2008) was used as an anatomical reference atlas for mouse brain structures and Bregma coordinates.

### Quantitative analysis of brain atrophy

A 2× objective was used to visualize brain atrophy following stroke. To measure brain atrophy, 2 sections per mouse, at Bregmas −1.70 mm and −1.94 mm were analyzed for anatomical abnormalities via Nissl staining. The area of the lateral ventricles and the thickness of the primary somatosensory cortex, from the corpus callosum through layers I-VI of the cortex of each hemisphere were manually traced for each section. The area of the ventricles and the thickness of the cortex were computed using NIH Image J software (Media Cybernetics, Rockville, MD).

### Quantitative analysis of neuronal degeneration

For quantitative analysis of of NeuN+ immunoreactivity, NeuN+ immunolabeled neurons in the primary somatosensory cortex adjacent to the infarct, and in the area of axonal degeneration in thalamic white matter regions were measured in 2 sections for each mouse. Section 1 was at the level of Bregma +1.18 mm, and section 2 was at the level Bregma +0.74 mm. Using a 10× objective, 2 fields were randomly selected from the ipsilateral cortex and thalamus for each section. Corresponding fields were taken from the contralateral hemisphere. For measurement of cholinergic neurodegeneration, ChAT+ immunolabeled cholinergic somas and their neurites in the medial septum of the basal forebrain were imaged. The total amount of ChAT+ immunoreactivity in the medial septum was quantified in 2 sections for each mouse with section 1 at Bregma +1.18 mm, and section 2 at Bregma +0.50 mm. For measurement of TH+ fiber projections, 2 fields were randomly selected from the primary somatosensory cortex adjacent to the infarct using a 10× objective. Three sections were analyzed per mouse for a total of 6 fields per mouse between Bregma +1.18 mm and Bregma +0.74 mm. Corresponding fields were taken from the contralateral hemisphere. The percent area occupied by NeuN+, ChAT+, and TH+ staining in each region was computed using NIH Image J software.

### Quantitative analysis of tau immunoreactivity

AT8/p-tau+ immunoreactivity in the degenerating white matter tracts of the thalamus were visualized with a 10× objective. In this analysis, 3 sections that contained the thalamus were selected; section 1 contained the thalamus at Bregma −0.70 mm, section 2 contained the thalamus at Bregma −1.46 mm, and section 3 contained the thalamus at Bregma −2.18 mm for each mouse. The total percentage occupied by p-tau+ staining in the thalamus on each section was computed using NIH Image J software. Sections from APP (amyloid precursor protein) (Nguyen et al., 2014; Nguyen et al., 2018) mutant mice displaying both amyloid and tau pathologies were used as a positive control for the pTau+ staining with the AT8 antibody.

### Quantitation of CD68, B220/CCD45R, and CD3e immunoreactivity in the stroke infarct

Immunoreactivity of CD68+ labeled microglia/macrophages, B220+ labeled B cells, and CD3e+ labeled T cells were visualized with a 10× objective and measured in the infarct using NIH Image J (Media Cybernetics, Rockville, MD). Three to 6 sections were stitched from 3-4 fields to capture the entire infarct area between Bregma −0.46 mm and Bregma −1.94 mm per mouse.

### Quantitation of CD68 and CD3e immunoreactivity in white matter tracts of the thalamus

CD68+ microglia and macrophages and CD3e+ T cells were visualized with a 10× objective and analyzed separately per mouse at Bregma −1.70 in the white matter tracts of the thalamus (area of axonal degeneration) following stroke surgery. Two non-overlapping, adjacent fields were randomly selected from 2 sections per mouse for CD68+ quantification from each hemisphere of the thalamus at the reticular thalamus nucleus to the ventral posteromedial thalamus nucleus. One field (2 sections per mouse for a total of 2 fields) was taken for CD3e+ quantification at the same brain region. The percentage occupied by CD68+ or CD3e+ immunostaining was computed.

### Ladder rung test

The ladder rung test was adapted from the ladder rung walking task (Rha et al., 2011; Doyle et al., 2012). The apparatus consisted of plexiglass rungs (10.16 cm length × 0.32 cm diameter) spaced at a constant distance of 0.64 cm apart, inserted between two plexiglass walls (0.64 cm width × 76.20 cm length × 15.24 cm height) and spaced 3.18 cm apart, providing a wide enough space for the animal to transverse the ladder. The apparatus was suspended 18 cm from the tabletop surface using blocks. The test was performed in a darkened room with a desk lamp illuminating the start zone to incentivize the animals to move across the ladder, and a small box was placed at the end zone. The mice were kept in a plastic chamber (9 cm × 8 cm × 14 cm) that was level with the plexiglass rungs at the start zone. As a mouse moved across the ladder, its steps were recorded from below, using a camera that slides along the surface of the testing table. Prior to pre-surgery testing, training was necessary to ensure that the animals were able to spontaneously traverse the apparatus. Eight training trials were required for animals to make an acceptable maximum number of limb placement errors (≤10-12%). Each mouse was tested once on each test day. The ladder apparatus was cleaned thoroughly with 10% ethanol and wiped dry between trials to minimize olfactory cues. Video recordings of each mouse traversing the ladder were analyzed at 0.25–0.30× playback speed using standard film playback software (VLC media player for Mac OSX). A trained observer focused on the front limb on the side of the body contralateral to the stroke infarct (i.e., front left paw). Once the mouse placed all four limbs on the ladder, the observer began counting correct steps and missteps of the front left paw. If the paw was placed on a rung and was not removed until the following step, a correct step was recorded. If the paw was placed on a rung and then slipped off the rung, or if the paw missed the rung entirely, a misstep was recorded. Two trained experimenters scored each video, and their values were averaged. The percent correct foot placement was calculated as 100 × (number of correct steps/number of correct steps + number of missteps).

### Nest construction test

Mice were transferred to individual testing cages with all environmental enrichment items (i.e., domes, tunnels) removed. One square (5 cm) piece of nestlet (Ancare, Bellmore, NY) made of pressed cotton batting, was placed in the same place of each test cage. Mice were provided with food and water. Testing was performed overnight, and images of nests built by the mice were captured the next morning. Nests were scored as follows on a scale of 1-5 based on how they were constructed: (i) nestlet was largely untouched (>90% remaining intact; 1 point); (ii) nestlet was partially torn up (50-90% remaining intact; 2 points); (iii) nestlet was mostly shredded, but often there was no identifiable nest site (<50% of the nestlet remaining intact and <90% within a quarter of the cage floor area, i.e., the cotton was not gathered into a nest, but scattered about the cage; 3 points); (iv) an identifiable, but flat nest (>90% of the nestlet shredded and the material was gathered into a nest within a quarter of the cage floor area, but the nest was flat, with walls higher than the mouse’s body height around <50% of its circumference; 4 points); and (v) a near perfect nest (>90% of the nestlet shredded and shaped into a crater, with walls higher than the mouse’s body height around >50% of its circumference; 5 points). This nest construction scoring system is based on Deacon and colleagues’ protocol (Deacon, 2006). Two trained experimenters scored each nest, and their values were averaged.

### Y-maze spontaneous alternation behavior (SAB) test

Mice were tested in a Y-shaped maze composed of beige acrylonitrile butadiene styrene (ABS) plastic and consisting of two symmetrical arms and one longer arm set at a 120° angle from each other (equal arms: 7.5 cm width × 37.0 cm length × 12.5 cm height; longer arm: 7.5 cm width × 42.0 cm length × 12.5 cm height). Mice were placed at the end of the longest arm of the maze, facing outward, and allowed to freely explore the 3 arms for 5 minutes. Over the course of multiple entries, mice normally exhibit a tendency to visit a new arm of the maze rather than visiting a recently visited arm. An entry was recorded when all four limbs of the mouse were within an arm of the maze. The number of arm entries and the number of triads were recorded and scored by the ANY-maze behavioral video tracking software (Stoelting, Wood Dale, IL), and the percentage of SAB was calculated by the software. The Y-maze apparatus was cleaned with 10% ethanol and wiped dry between trials to minimize olfactory cues.

### Novel object recognition (NOR) test

The NOR arena consisted of a white opaque box measuring 38.0 cm width × 48.0 cm length × 20.0 cm height made of ABS plastic. Mice were habituated to the NOR arena over the course of 2 days prior to testing day. During the first day of habituation, mice were placed with cage mates for 15 minutes in the NOR arena. The following day, each mouse was allowed to explore the arena alone for 15 minutes. On testing day, each mouse first underwent an “object familiarization phase,” during which they were placed in the arena along with two identical objects located at different corners of the arena 5 cm from the walls. The objects consisted of children’s building blocks of different shapes and colors. Mice were habituated to the NOR arena over the course of 2 days prior to testing day. During the first day of habituation, mice were placed with cage mates for 15 minutes in the NOR arena. The following day, each mouse was allowed to explore the arena alone for 15 minutes. On testing day, each mouse first underwent an “object familiarization phase”, during which they were Published studies confirmed no innate preference between objects. Mice were allowed to explore the arena and objects for 5 minutes. Object exploration was defined as contact with the object by the mouse’s nose within 2 cm of the object, which was recorded by ANY-maze behavioral video tracking software 4 hours following the object familiarization phase, each mouse underwent an “object recognition testing phase”, during which they were placed back in the arena with one of the objects they previously explored during the familiarization phase [the familiar object (FO)], and a novel object (NO). The object role (novel versus familiar) and position (left versus right) were balanced within each experimental group. Different sets of objects were used for each timepoint to control for biases from previously exposed objects. ANY-maze behavioral video tracking software was used to compute the time each mouse spent exploring each object. The recognition index (RI) was calculated as (NO Time)/(NO Time + FO Time) and expressed as a percentage by multiplying by 100. The NOR apparatus was cleaned with 10% ethanol between trials to minimize olfactory cues.

### Open field (OF) testing

The OF test assesses exploratory habits in a relatively large novel environment. The OF apparatus consisted of a beige ABS plastic arena measuring 50 cm length × 50 cm width × 50 cm height. The arena was divided into 10 cm squares (grid). The center zone of the arena measured 30 cm length × 30 cm width. The periphery zone included all the squares along the edge of the arena and measured 8 cm width. The space between the center and periphery zones was considered the “dead” zone. Each mouse was placed into the arena and allowed to explore for 5 minutes. During these 5 minutes, ANY-maze software was used to track the center of the animal’s body within the arena. The software then computed the number of entries and exits into each zone for each mouse. The OF apparatus was cleaned with 10% ethanol and wiped dry between trials to minimize olfactory cues.

### Power analysis and sample size calculations

Sample sizes were determined by a priori power analysis with data from our pilot studies and previous published studies (Nguyen et al., 2014; Nguyen et al., 2016; Nguyen et al., 2018). For the metabolomics and multiplex immunoassay analyses (Phase I of the study), according to the calculations, the minimum number of mice needed to attain statical significance of p<0.05 with an 80% probability was n=5 mice/experimental group when we anticipated a ≥20% difference between experimental groups. For the behavioral studies (Phase II), the minimum number of mice needed to attain statistical significance of p<0.05 with an 80% probability was n=9 mice/experimental group when we anticipated a ≥20% difference between experimental groups. Therefore, we included a total of 10 male mice in each experimental group to account for possible deaths due to age or surgery for behavior tests. All the mice used for behavior tests were also used for morphological studies (Phase II of the study), even though the minimum number of mice needed to attain statistical significance of p<0.05 with an 80% probability was n=5 mice/experimental group when we anticipated a ≥20% difference between experimental groups for this analysis.

### Statistics

Data are presented as mean ± SD. Datasets containing 3 groups were analyzed using a one-way ANOVA, followed by *post-hoc* Newman-Keuls testing. Datasets containing 2 groups were evaluated by Student’s t-test. Significance was set at p<0.05. For all figures, *p is <0.05, **p is <0.01, ***p is <0.001, and ****p is <0.0001. The statistical summary of the metabolomics data is shown in Table 1. Metabolon’s proprietary Pathway Surveyor tool was used to identify key pathways significantly altered by stroke and LM11A-31 treatment.

**Table 1.**
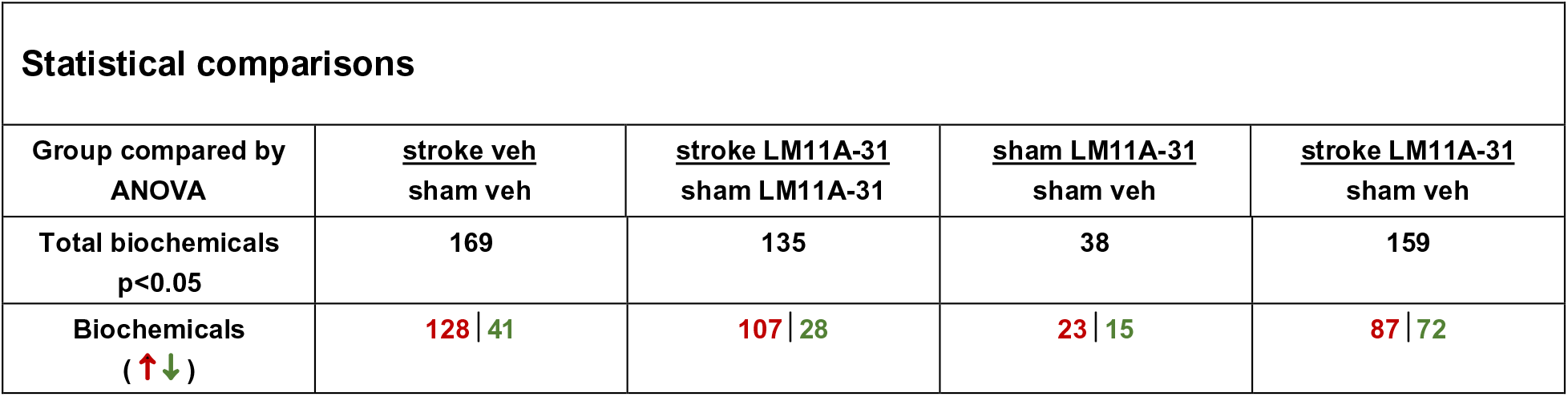
Total numbers of biochemicals that significantly differed (p≤0.05) between the listed experimental groups after global metabolic profiling from a total of 645 biochemicals analyzed.

| Statistical comparisons |  |  |  |  |
| --- | --- | --- | --- | --- |
| Group compared by ANOVA | <u>stroke veh</u><br>sham veh | <u>stroke LM11A-31</u><br>sham LM11A-31 | <u>sham LM11A-31</u><br>sham veh | <u>stroke LM11A-31</u><br>sham veh |
| Total biochemicals<br>p<0.05 | 169 | 135 | 38 | 159 |
| Biochemicals<br>(↑↓) | 128 \| 41 | 107 \| 28 | 23 \| 15 | 87 \| 72 |

## RESULTS

### Study design

The goal of this study was to evaluate the therapeutic effect of a p75^NTR^ small molecule ligand on recovery from stroke. To achieve this goal, the study was divided into two testing phases. For Phase I (Figure 1A, Study design), mice were randomized and underwent stroke or sham surgery. A week after surgery, mice were orally administered LM11A-31 for 7 days per week, one dose per day, for 2 weeks, beginning 1 week after surgery (experimental groups: n=5 for sham-veh, n=5 for stroke-veh, and n=6 for stroke-31). At 3 weeks following surgery, mice were euthanized, and their infarcts were processed for multiplex immunoassay. The remainder of their ipsilateral hemispheres was processed for metabolomics. The goal of Phase I was to evaluate how LM11A-31 alters brain biochemistry and post-stroke inflammation during treatment.

**Figure 1.**
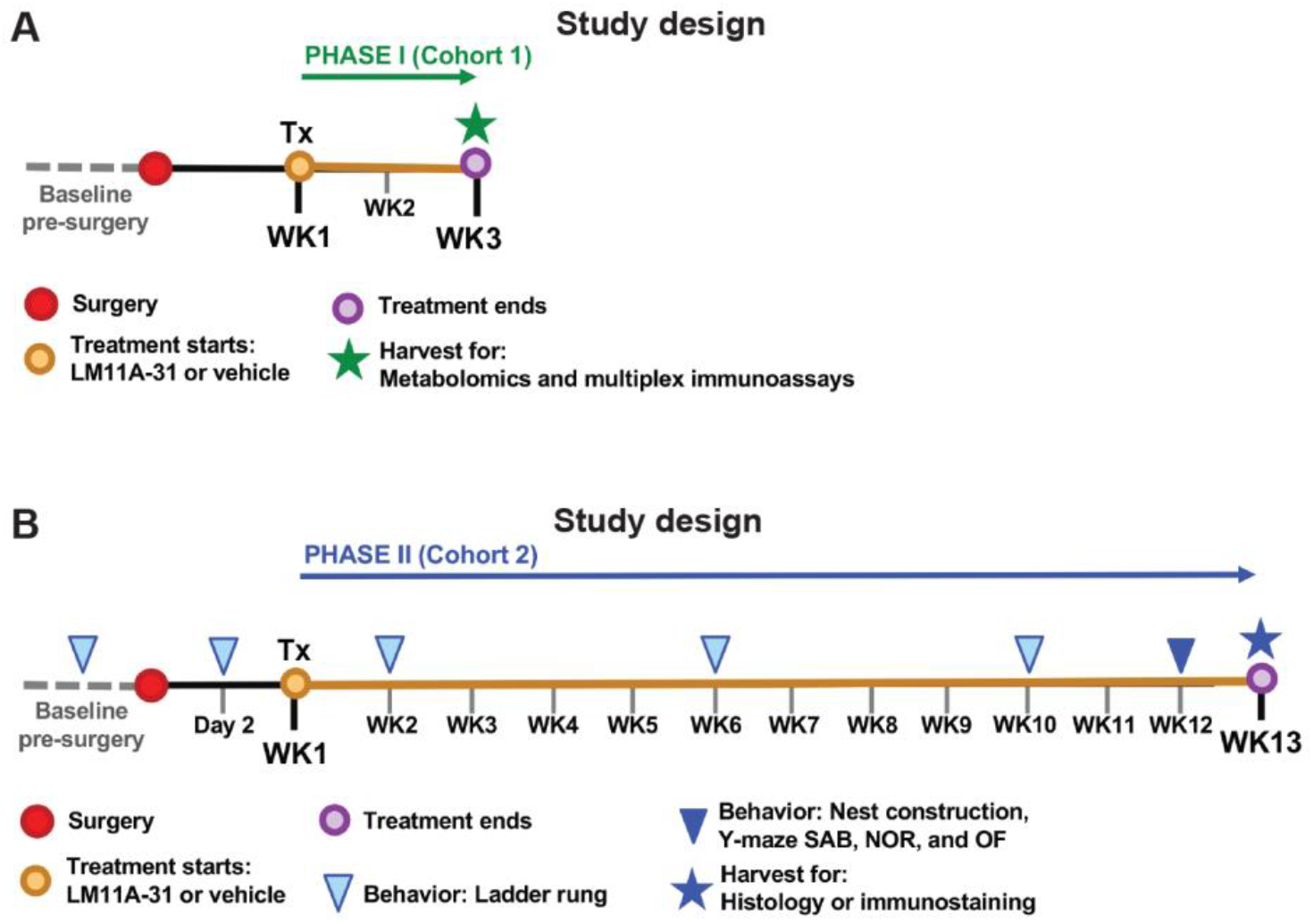
Study design for oral administration of 50 mg/kg LM11A-31 to aged mice as a treatment for stroke. (**A**) For Phase I (Cohort 1), mice were administered (Tx: treatment) LM11A-31 or vehicle one time per day for a total of 2 weeks, beginning 1 week following stroke or sham surgery. At 3 weeks following surgery the infarct from each mouse was harvested for multiplex immunoassay. The remainder of each ipsilateral hemisphere was harvested for metabolomics. (**B**) For Phase II (Cohort 2), stroked mice were administered (Tx: treatment) LM11A-31 or vehicle 5 days per week for a total of 12 weeks, beginning 1 week following surgery. At 13 weeks following stroke surgery, brains were harvested for histology and immunostaining. In Phase II, mice also underwent behavioral testing at the indicated timepoints.

For Phase II (Figure 1B, Study design), a separate cohort of mice underwent baseline ladder rung testing, and their scores were used to stratify mice into either LM11A-31 or vehicle treatment groups (experimental groups: n=10 for stroke-veh, and n=10 for stroke-31). Mice then underwent stroke surgery and were administered with either LM11A-31 or vehicle for 5 days per week, one dose per day, by oral administration for 12 weeks, beginning 1 week after surgery. The ladder rung test was repeated at 2 days, 2 weeks, 6 weeks, and 10 weeks after stroke surgery. During the 12th week of treatment, mice were tested on a panel of behavioral tests, including a nest construction test, a Y-maze Spontaneous Alternation Behavior (SAB) test, a Novel Object Recognition (NOR) test, and an Open Field (OF) test. After 12 weeks of treatment (13 weeks post-stroke), mice were euthanized, and their brains were processed for histology and immunostaining. The goal of Phase II was to evaluate treatment efficacy of LM11A-31 in the context of stroke recovery.

### Phase I: Global metabolic profiling of LM11A-31 treated mice

After 2 weeks of treatment (3 weeks post-stroke or sham surgery), global metabolic profiling of the ipsilateral hemispheres from the mice treated with LM11A-31 and vehicle returned measurements for 645 compounds. Following log transformation and imputation of missing values with the minimum observed value for each compound, a three-way ANOVA was performed to identify the chemicals that were significantly different between experimental groups.

Among the mice that were treated with vehicle alone, 169 chemicals were present at different signal intensities in stroked mice compared to sham operated mice. Among the mice that were treated with LM11A-31, 135 chemicals were present at different intensities in stroked compared to sham operated mice. The least difference in metabolic profile, which equated to changes in 38 chemicals, was detected among mice that underwent sham surgery, between those treated with LM11A-31 and those treated with vehicle. However, the equivalent comparison among stroked mice showed 159 chemicals that were altered by treatment with LM11A-31. These data indicate that (i) there are substantial alterations in brain metabolism at 3 weeks following stroke; (ii) that treatment with LM11A-31 in sham operated mice does not substantially alter brain metabolism; and (iii) that 2 weeks of LM11A-31 versus vehicle administration substantially alters chronic changes in brain metabolism induced by stroke.

Principle component analysis (PCA) was used to visually assess the separation between the experimental groups (Figure 2). There was both a separation along Component 1 due to treatment (Figure 2B **&** C), and along Component 2 due to stroke (Figure 2B **&** D). This analysis demonstrated that there was sub-clustering based on both compound treatment and disease state. A summary of the numbers of chemicals that achieved statistical significance (p≤0.05), as well as whether they were increased or decreased is shown in Table 1, and a full list of the metabolic changes are provided in **Supplementary Tables 1 and 2**.

**Figure 2.**
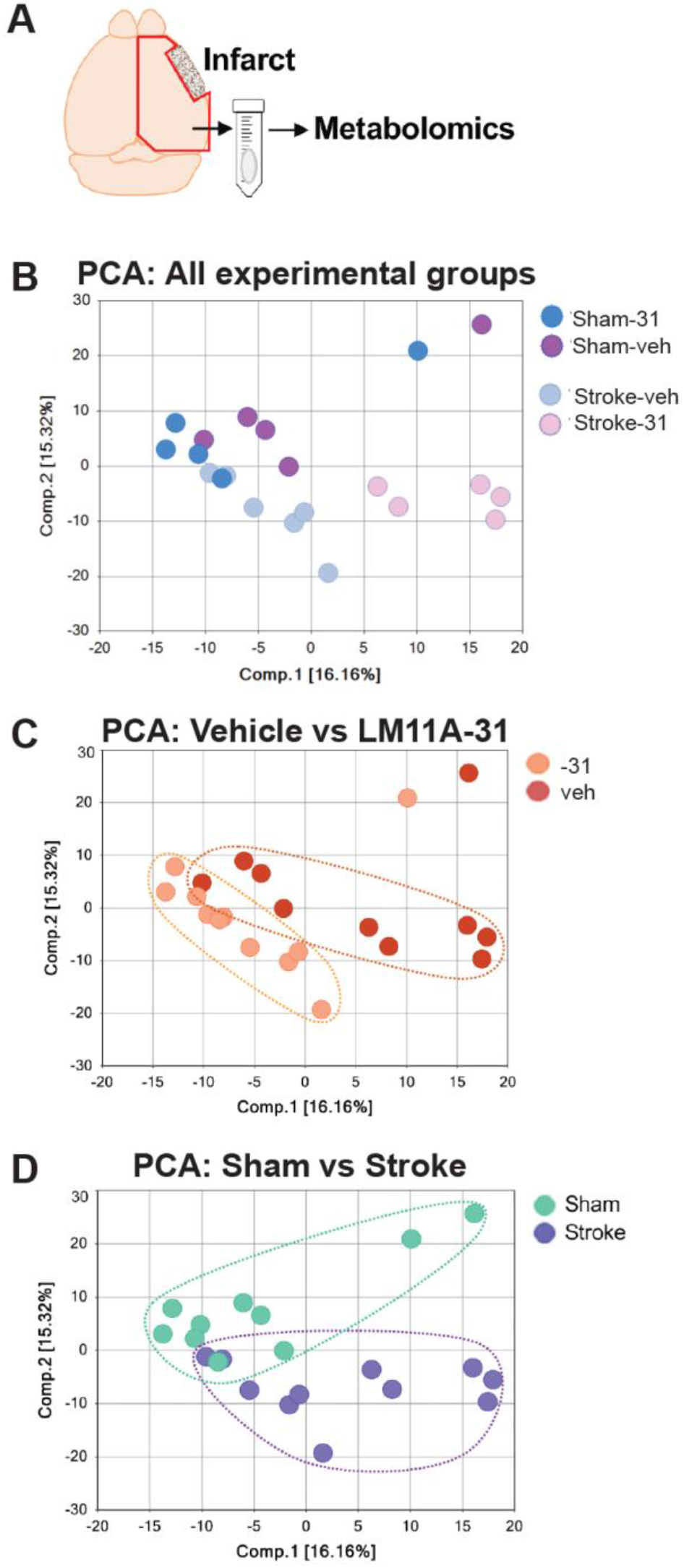
Principal component analysis (PCA) to allow for visual assessment of the clustering of each sample within each group. (**A**) Area dissected for global metabolic profiling. (**B**) PCA analysis with all four groups represented. (**C**) PCA reveals separation between vehicle and LM11A-31 treated mice along component 1 demonstrating a strong drug treatment effect. (**D**) PCA reveals separation between mice that were subjected to sham and stroke surgery along component 2, demonstrating a strong disease effect.

Next, we used a pathway surveyor tool to evaluate which key pathways are altered by stroke and by LM11A-31 treatment after stroke. This finding revealed that LM11A-31 can mitigate stroke induced decreases in the levels of multiple neurotransmitters, as well as stroke induced changes in redox homeostasis and glucose metabolism. With regard to neurotransmitter levels, we found that following stroke, mice had lower levels of glutamate, serotonin, and acetylcholine in the ipsilateral hemisphere compared to mice that underwent sham surgery (Figure 3A-C). Treatment with LM11A-31 after stroke significantly increased the levels of each of these neurotransmitters, except for glutamate. Notably, levels of dopamine trended towards a significant reduction following stroke (p=0.1038) and were also significantly increased by LM11A-31 treatment (Figure 3D).

**Figure 3.**
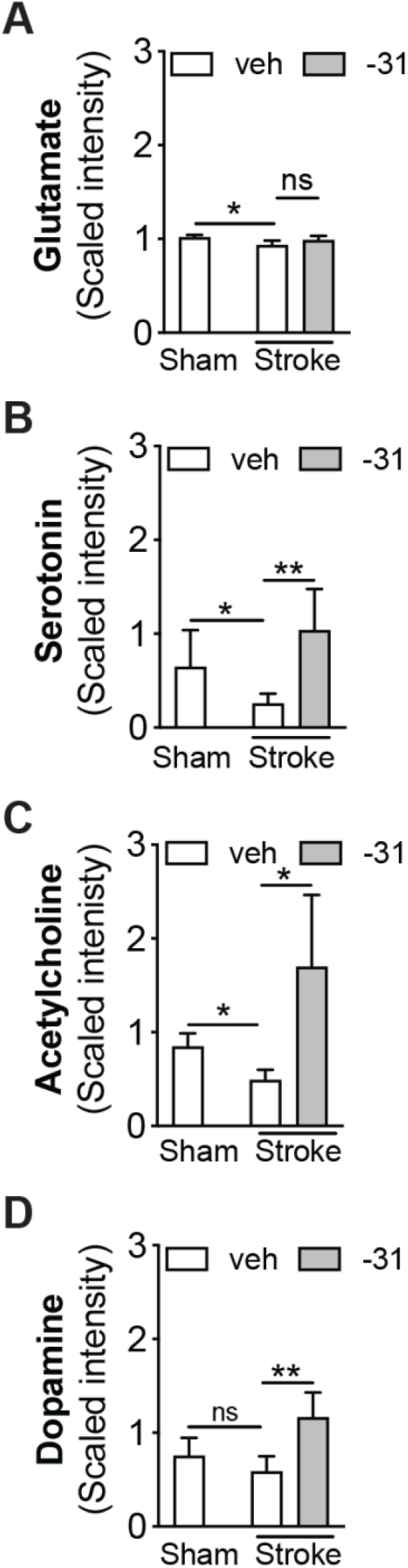
LM11A-31 attenuated stroke induced reductions in glutamate, serotonin, and acetylcholine, and increased levels of dopamine. Three weeks following stroke, the levels of **(A)** glutamate, **(B)** serotonin, and **(C)** acetylcholine were significantly reduced in the ipsilateral hemisphere compared to sham operated mice. The reductions in serotonin and acetylcholine were attenuated following 2 weeks of LM11A-31 (−31) compared to vehicle (veh) treatment. (**D**) Dopamine was not significantly reduced following stroke; however, it was significantly increased by 2 weeks of LM11A-31 compared to vehicle treatment. Group sizes: n=5 sham-veh, n=5 stroke-veh, n=6 stroke-31. Data represent mean ± SD. *p<0.05 and **p<0.01.

Regarding redox homeostasis, global metabolic profiling revealed that stroke compared to sham surgery resulted in a decrease in reduced glutathione (GSH) in the ipsilateral hemisphere. LM11A-31 restored GSH in stroked mice (Figure 4A). LM11A-31 also increased levels of oxidized glutathione (GSSG; Figure 4B). These results indicate that LM11A-31 treatment increases the pool of GSH available to modulate chronic changes in redox homeostasis caused by stroke.

**Figure 4.**
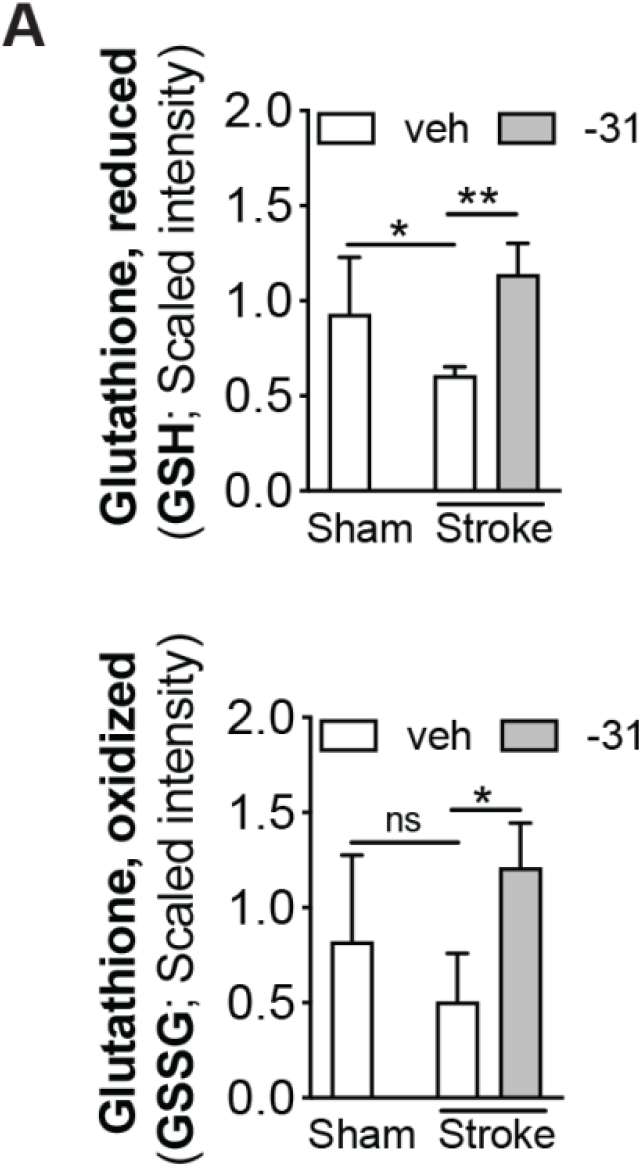
LM11A-31 altered redox homeostasis after stroke. (**A**) Compared to sham operated mice, stroked mice had significantly lower levels of reduced glutathione (GSH) in the infarcted hemisphere. Stroked mice treated with LM11A-31 (−31) for 2 weeks had a significantly higher levels of GSH than stroked mice treated with vehicle (veh) alone, suggesting a prevention or reversal of the stroke induced decrease in GSH. (**B**) Levels of oxidized glutathione (GSSG) were also increased in LM11A-31-treated mice compared to vehicle treated mice. The increase in GSSG suggests there is a larger pool of GSH available to mitigate oxidative stress caused by the chronic sequelae of stroke. Group sizes: n=5 sham-veh, n=5 stroke-veh, n=6 stroke-31. Data represent mean ± SD. *p<0.05, and **p<0.01.

Regarding glucose homeostasis (glycolysis pathway shown in Figure 5A), although levels of glucose were undetected, levels of pyruvate were reduced in the ipsilateral hemisphere of the mice that underwent stroke compared to sham surgery (Figure 5B). This finding indicates that there is a reduction in glycolysis in the ipsilateral hemisphere at 3 weeks after stroke. There was also a trend towards a reduction in acetyl-CoA, the oxidative decarboxylation product of pyruvate (Figure 5B). This finding is an indication that the reduction in glycolysis reduces the amount of acetyl-CoA available for the tricarboxylic acid (TCA) cycle. Treatment with LM11A-31 after stroke increased levels of acetyl-CoA (Figure 5B).

**Figure 5.**
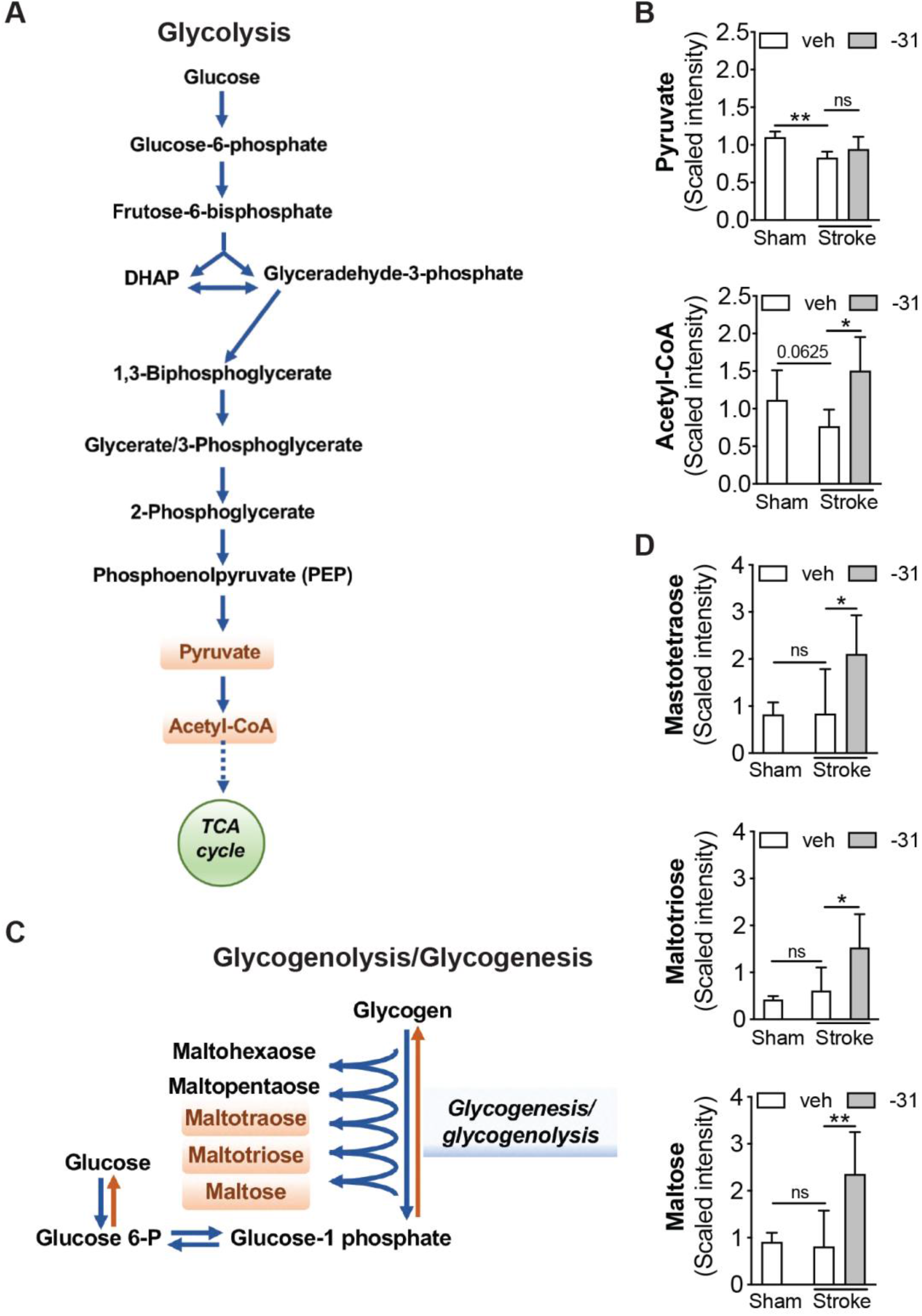
LM11A-31 altered glucose metabolism after stroke. (**A**) The glucose metabolism pathway or glycolysis. (**B**) Compared to sham operated mice, stroked mice had significantly lower levels of pyruvate (*top graph*) in their brains and showed a trend towards lower levels of acetyl-CoA (p=0.0625; *bottom graph*). Although 2 weeks of LM11A-31 (−31) treatment did not significantly increase pyruvate levels compared to vehicle (veh) treatment, it did significantly increase levels of acetyl-CoA compared to vehicle treatment. (**C**) Glycogenolysis/Glycogenesis. (**D**) Although not altered by stroke, maltotetraose (*top graph*), maltotriose (*middle graph*), and maltose (*bottom graph*) were all increased following LM11A-31 treatment. This suggests that an increase in glycogenolysis may be the mechanism by which LM11A-31 increased acetyl-CoA after stroke. Group sizes: n=5 sham-veh, n=5 stroke-veh, n=6 stroke-31. Data represent mean ± SD. *p<0.05 and **p<0.01.

Treatment with LM11A-31 after stroke also altered the glycogenesis/glycogenolysis pathway (Figure 5C). This finding is supported by increases in maltotetraose, maltotriose, and maltose in the ipsilateral hemispheres of the LM11A-31 treated mice compared to the vehicle treated mice after stroke (Figure 5D). Like glucose, glycogen was undetected and so it is unclear whether these differences in intermediates are the result of increased glycogenolysis or increased glycogenesis. However, combined with the elevated level of acetyl-CoA seen in LM11A-31 treated mice compared to vehicle treated mice following stroke, these findings provide evidence that LM11A-31 treatment increases glycogenolysis, thereby increasing the amount of glucose available for acetyl-CoA generation.

Acetyl-CoA can also be generated by beta oxidation of long-chain fatty acids. To determine whether the increase in acetyl-CoA was the result of increased fatty acid oxidation we evaluated whether LM11A-31 treatment after stroke increased the levels of carnitine and acetyl-carnitine, key metabolites required for the generation of acetyl-CoA by beta oxidation. Although levels of carnitine and acetyl-carnitine were increased by stroke, they were not further increased by LM11A-31 treatment after stroke (**Supplementary Tables 1 and 2**). This result indicates that the increase in acetyl-CoA levels in response to LM11A-31 treatment after stroke is unlikely due to an increase in the beta oxidation of long-chain fatty acids.

### Phase I: Evaluation of cytokine/chemokine levels in the infarct in LM11A-31 treated mice

We have previously provided evidence that stroke infarcts are sites of chronic inflammation (Doyle et al., 2015; Nguyen et al., 2016; Zbesko et al., 2018). To determine whether LM11A-31 has an effect on chronic inflammation in the infarct, we used a multiplex immunoassay to measure cytokines/chemokines within the infarcts of Phase I mice and tissue from the equivalent region of the cortex from sham operated mice. Analytes of interest included granulocyte colony-stimulating factor (G-CSF), granulocyte-macrophage colony-stimulating factor (GM-CSF), interferon-γ (IFN-γ), interleukin (IL)-1α, IL-1Β, IL-2, IL-4, IL-5, IL-6, IL-7, IL-9, IL-10, IL-12(p40), IL-12(p70), IL-13, IL-15, IL-17, interferon–induced protein-10 (IP-10), keratinocytes-derived chemokine (KC), monocyte chemoattractant protein-1 (MCP-1), macrophage inflammatory protein-1α (MIP-1α), macrophage inflammatory protein 1β (MIP-1β), macrophage inflammatory protein-2α (MIP-2α), regulated on activation normal T cell expressed and secreted (RANTES), and tumor necrosis factor-α (TNF-α). Out of these 25 cytokines/chemokines, only MIP-2α, IP-10, RANTES, and MIP-1β were increased in the infarct site at 3 weeks after stroke when compared to tissue from the equivalent hemisphere of sham operated mice. LM11A-31 did not alter the levels of these cytokines/chemokines (Figure 6). These data indicate that two weeks of LM11A-31 treatment does not ameliorate or exacerbate inflammation within the infarct site at 3 weeks following stroke.

**Figure 6.**
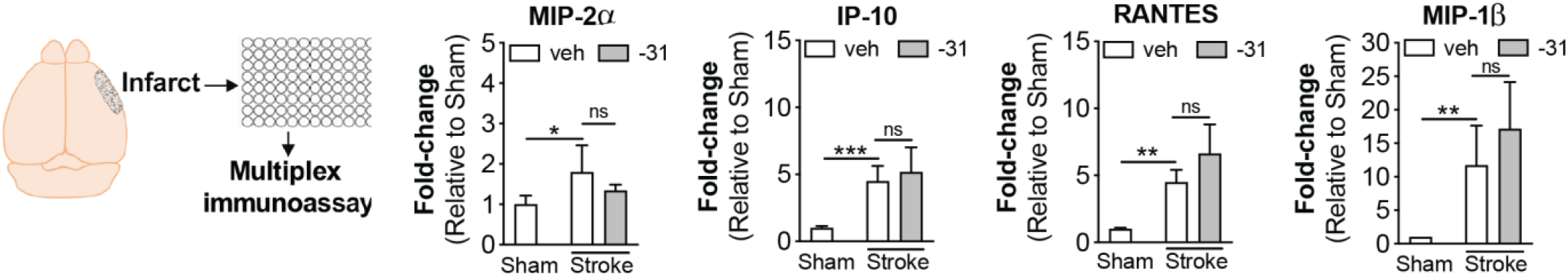
LM11A-31 did not alter cytokine levels in the infarct. (**A**) Out of 25 cytokines measured by multiplex immunoassay, only MIP-2α, IP-10, RANTES, and MIP-1β were significantly increased in the infarct at 3 weeks following stroke compared to sham surgery. Treatment with LM11A-31 for 2 weeks, beginning 1 week after stroke did not significantly alter the levels of these cytokines. Group sizes: n=5 sham-veh, n=5 stroke-veh, n=6 stroke-31. Data represent mean ± SD. *p<0.05, **p<0.01, and ***p<0.001.

### Phase II: Evaluation of LM11A-31 on peripheral immune cell infiltration in the infarct and area of axonal degeneration

The goal of Phase II was to evaluate the efficacy of LM11A-31 at ameliorating chronic neuropathology and neurodegeneration when treatment is initiated one week after stroke and after 12 weeks of daily dosing. The impact of LM11A-31 on inflammation in the infarct was determined by immunostaining of CD68+ microglia/macrophages, B220+ B cells, and CD3e+ T cells at the site of the infarct. Percent area occupied by CD68+, B220+, and CD3e+ immunostaining revealed that LM11A-31 treatment after stroke did not alter macrophage, B lymphocyte or T lymphocyte cell infiltration into the infarct at 13 weeks (Figure 7A-C).

**Figure 7.**
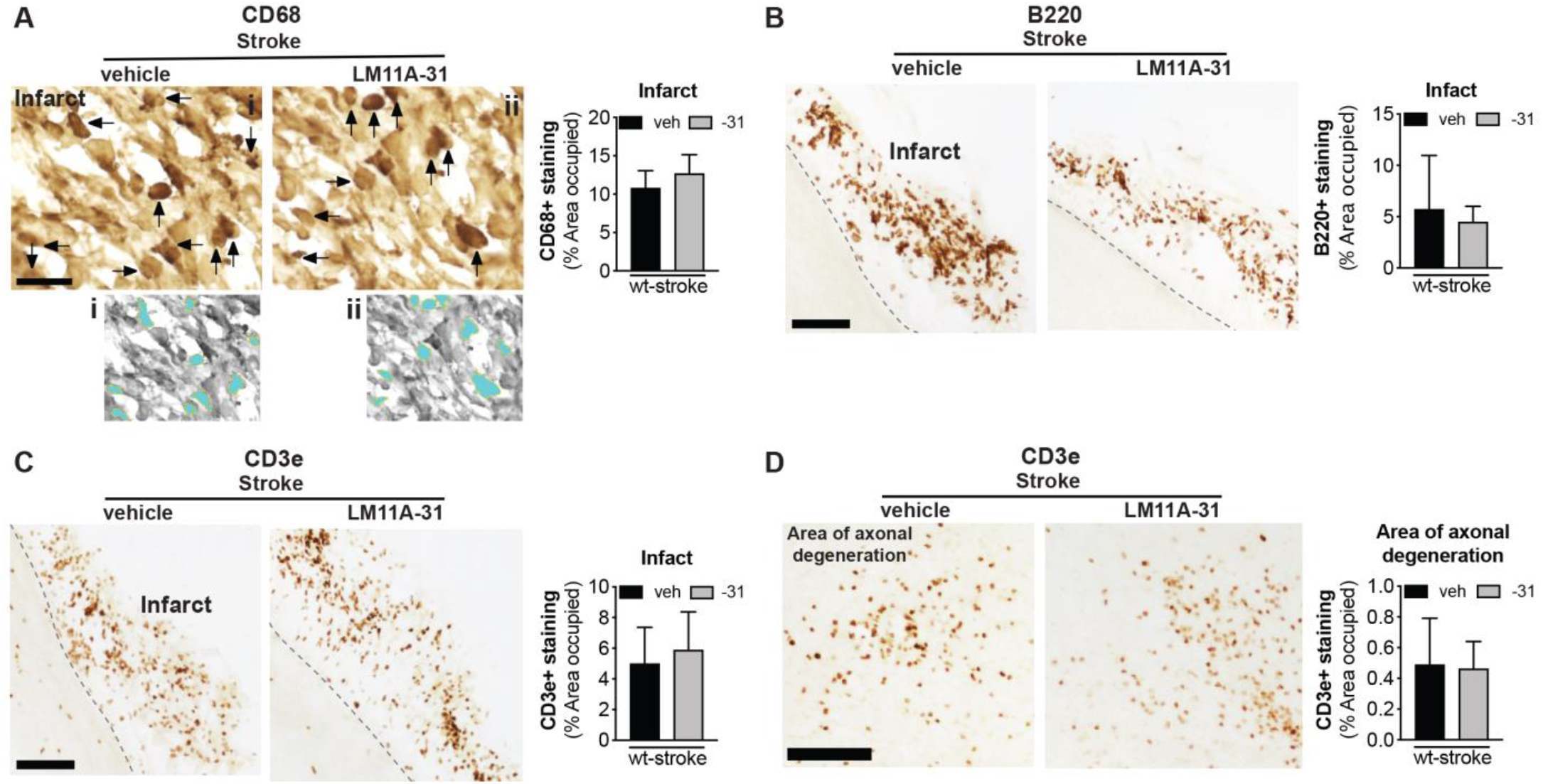
LM11A-31 did not alter immune cell infiltration in the infarct or T cell infiltration in the area of axonal degeneration. (**A**) Representative images and quantification of CD68 immunolabeled microglia and macrophages in the infarcts of mice given vehicle (veh) or LM11A-31 (−31) for 12 weeks, starting 1 week post-stroke. Panels (i) and (ii) show the method of thresholding used to measure the % area covered by cellular CD68+ immunostaining. Stroked mice given vehicle and stroked mice given LM11A-31 displayed similar CD68+ immunoreactivity; quantification revealed no significant differences between these treatment groups. (**B**) B220+ immunolabeled B cells and (**C**) CD3e+ immunolabeled T cells in the infarcts of mice given vehicle or LM11A-31 for 12 weeks, starting 1 week post-stroke. Stroked mice given vehicle and stroked mice given LM11A-31 also displayed similar B220+, and CD3e+ immunoreactivity; quantification revealed no significant differences between mice treated with either agent. (**D**) Representative images of CD3e+ immunolabeled T cells in white matter tracts projecting from the infarct area. There was no difference in CD3e+ immunoreactivity in stroked mice given vehicle or LM11A-31 in this area of axonal degeneration. Scale bars, 125 μm. Group sizes: n=10 stroke-veh, n=10 stroke-31. Data represent mean ± SD.

We previously reported that in the weeks after stroke, CD3e+ T cells are not only present in the infarct, but are also present in the thalamus, striatum, and internal capsule in the ipsilateral hemisphere (Doyle et al., 2015). These are locations of degenerating axons that project from the sensorimotor cortex, which is the cortical region infarcted in the DH stroke model. Treatment with LM11A-31 did not affect T cell infiltration into the thalamic white matter tracts of the area of axonal degeneration (Figure 7D).

### Phase II: Evaluation of the impact of LM11A-31 on secondary neurodegeneration

Stroke causes delayed ipsilateral cortical atrophy and tissue thinning in 3-5-month-old C57BL/6 mice at 8 weeks post-stroke (Nguyen et al., 2018). This atrophy is more pronounced and results in enlargement of the ipsilateral lateral ventricle in 18-month-old mice (Nguyen et al., 2018). Therefore, we next sought to determine whether brain atrophy caused by stroke can be prevented or reduced by treatment with LM11A-31. As shown in the representative images, stroked mice that were treated with LM11A-31 demonstrated a significant reduction in the degree of post-stroke ventricular enlargement compared to those treated with vehicle (Figure 8A). Additionally, vehicle treated mice had significantly thinner ipsilateral primary somatosensory cortices compared to the contralateral cortices, and the equivalent cortices of LM11A-31-treated mice following stroke (Figure 8B). These data demonstrate that LM11A-31 can ameliorate atrophy of the ipsilateral hemisphere following stroke.

**Figure 8.**
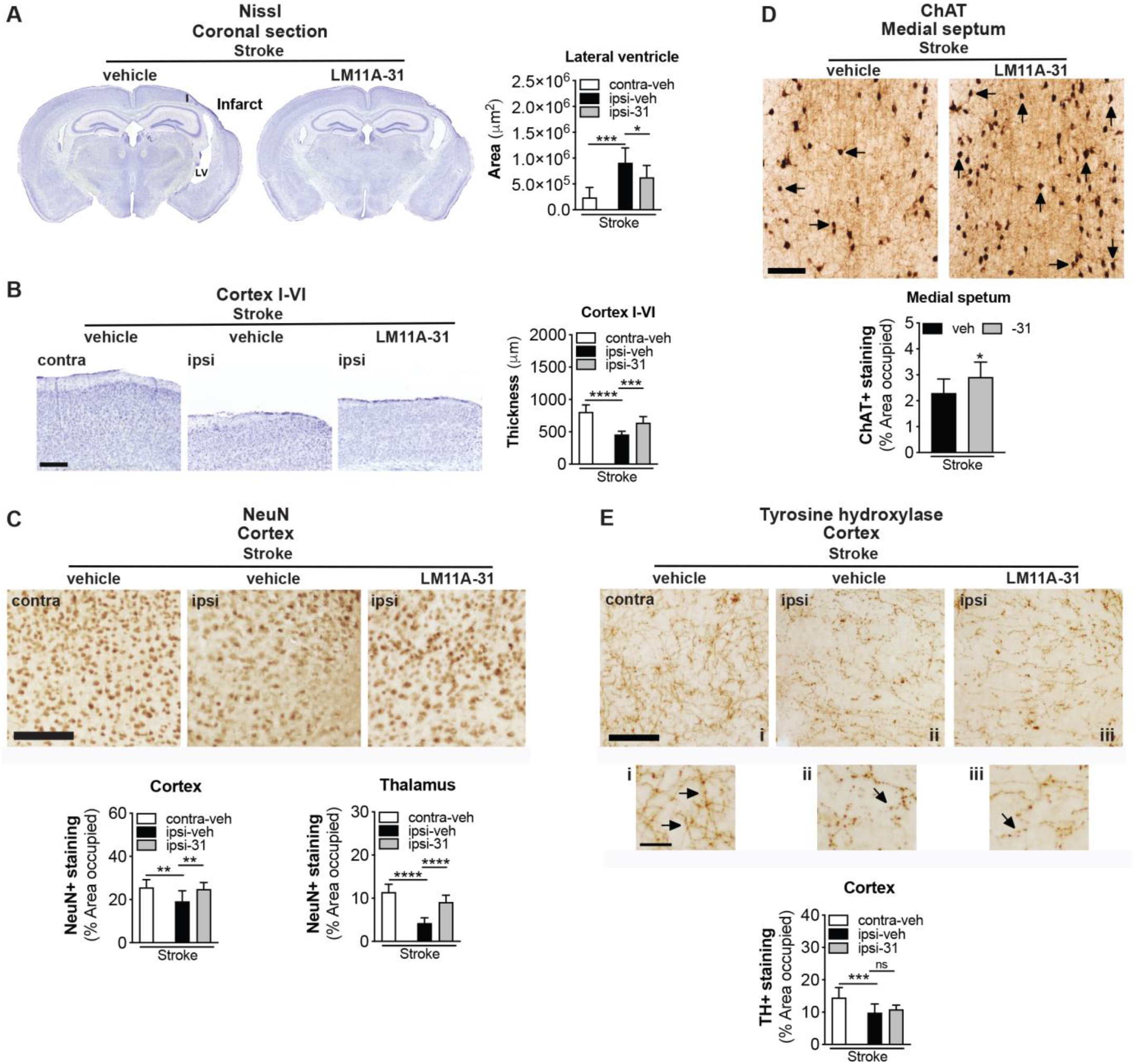
LM11A-31 reduces brain atrophy and preserves NeuN and ChAT immunoreactivity following stroke. (**A**) Representative 2× images of Nissl stained whole brain sections from mice given vehicle (veh) or LM11A-31 (−31) for 12 weeks, starting 1 week post-stroke (*left images*). Quantification of the lateral ventricle in the ipsilateral (ipsi) hemisphere revealed significant ventricle enlargement in stroked mice, which LM11A-31 significantly reduced (*right graph*). (**B**) Representative 2× images of cortical layers I-VI of the primary somatosensory cortex contralateral (contra) and ipsilateral (ipsi) to the infarct (*left images*). Quantification of cortical thickness in the ipsilateral compared to contralateral hemisphere revealed significan t tissue thinness in stroked mice. Stroked mice treated with LM11A-31 showed significantly thicker tissue in this area than stroked mice treated with vehicle alone (*right graph*). Scale bar, 250 μm (**C**) Representative 10× images of neuronal nuclei (NeuN) labeled in the ipsilateral primary somatosensory cortex (*top images*). Scale bar, 125 μm. Quantification revealed a significant reduction of NeuN+ staining in stroked mice, which was mitigated by LM11A-31 treatment (*middle and bottom graphs*). (**D**) Representative 10× images of choline acetyltransferase (ChAT)+ somas and neurites, and innervating projection fibers in the medial septum (*top images*). Many neurites (arrows) in the vehicle treated stroked mice displayed qualitative degenerative changes, including decreased length, which was not as evident in mice treated with LM11A-31. Scale bar, 125 μm. Quantification of cholinergic neurons, neurites, and fiber projections revealed a significant reduction of ChAT+ staining in stroked mice, which LM11A-31 also mitigated (*bottom graph*). (**E**) Representative 10× images of tyrosine hydroxylase (TH)+ projection fibers in the primary somatosensory cortex (*top images and magnified outsets i-iii*). Compared to sham operated mice, stroked mice displayed less abundant and more broken or disconnected fibers, which LM11A-31 treatment did not affect. Scale bar, 125 μm (40 μm for outsets). TH+ fiber quantification revealed significantly less TH+ immunostaining in stroked mice compared to sham operated mice, which was not altered by LM11A-31 treatment (*bottom graph*). Group sizes: n=10 stroke-veh, n=10 stroke-31. Data represent mean ± SD. *p<0.05, **p<0.01, ***p<0.001, and ****p<0.0001.

Stroke induced cortical atrophy correlates with neurodegeneration, as demonstrated by a reduction of neuronal nuclei marker, NeuN+, in the peri-infarct cortex of mice following stroke (Chung et al., 2018; Zbesko et al., 2018). As depicted in the representative images and quantification of stroked mice treated with LM11A-31 compared to vehicle, LM11A-31 treatment attenuated this reduction in NeuN+ immunoreactivity in the ipsilateral hemisphere (Figure 8C, *top images*). Specifically, there was a reduction in NeuN+ expression in the ipsilateral compared to the contralateral hemisphere of the stroked mice, and LM11A-31 administration resulted in a preservation of NeuN+ immunoreactivity (Figure 8C, *bottom graphs*). These data demonstrate that LM11A-31 can decrease the loss of NeuN+ immunoreactivity that occurs in the ipsilateral hemisphere in aged mice following stroke.

Most of the cerebral cortex, such as the prefrontal and the motor cortices, along with the somatosensory cortex, are connected to basal forebrain cholinergic neurons (Wu et al., 2014; Gielow and Zaborszky, 2017). Brain regions with synaptic connections to ischemic regions can undergo transneuronal degeneration after stroke (Zhang et al., 2012; Chang et al., 2016), and previously we observed approximately 30% less ChAT+ staining in the medial septum of the basal forebrain of 18-month-old C57BL/6 mice at 8 weeks post-stroke compared to sham surgery (Nguyen et al., 2018). In the present study, there was significantly more ChAT+ immunoreactivity in the medial septum of the mice treated with LM11A-31 compared to mice administered vehicle (Figure 8D).

Though primarily expressed in basal forebrain cholinergic neurons and pyramidal cortical and hippocampal neurons (Mufson EJ et al. 1992 PNAS 89:569-573; Miller MW and et al. 2000 Brain Res 852:355-366; Dougherty KD et al. 1999 J Neurosci 19:10140-10152; Zagrebelsky M et al. 2005 J Neurosci 25:9989-9999), p75^NTR^ is also present in neurons of the raphe nuclei, reticular formation, and locus coeruleus (noradrenergic neurons that contribute to the regulation of mood and behavioral states). TH+ neurites of noradrenergic neurons in the rostral portion of the locus coeruleus innervate hippocampal and cortical areas, and are markedly affected in, and correlate with, severity of cognitive decline (Grudzien et al., 2007) in AD and other neurodegenerative diseases, such as Down syndrome, Parkinson’s disease, and Huntington’s disease (Iversen et al., 1983; Mann, 1988; Zweig et al., 1992; Zarow et al., 2003). To determine whether stroke results in a loss of TH+ terminals innervating the peri-infarct cortical region, the density of TH+ labeled projections fibers in the ipsilateral and contralateral cortex was quantified. As shown in Figure 8E (*top images*), TH+ projection fibers appeared diminished in density within the ipsilateral primary somatosensory cortex, adjacent to the infarct, compared to the contralateral cortex in the stroked mice. There was a significant decrease in TH+ expression in the ipsilateral compared to the contralateral hemisphere of the stroked mice and LM11A-31 administration did not attenuate this reduction (Figure 8E, *bottom graph*). Together, these data provide evidence that LM11A-31 can reduce brain atrophy and preserve NeuN+ and ChAT+ immunoreactivity following stroke but is less effective at preventing the loss of TH+ projection fibers.

### Phase II: Evaluation of the effect of LM11A-31 on microglial activation and tau hyperphosphorylation

Ser202 and Thr205, the epitopes recognized by the antibody AT8 when they are phosphorylated, are among the tau residues that contribute to the pathological conformations of p-tau (Jeganathan et al., 2008). Previously, we reported that 18-month-old stroked mice develop tau pathology in areas of degenerating white matter tracts following stroke (Nguyen et al., 2018). Furthermore, we have shown that excessive levels of p-tau+ staining are reduced in APP mutant mice after LM11A-31 treatment (Nguyen et al., 2014). Consequently, we determined whether LM11A-31 reduces p-tau accumulation in degenerating white matter tracts following stroke. As seen in Figure 9A (*top images*), quantitation revealed that (i) the ipsilateral hemisphere contained more p-tau deposits than the contralateral hemisphere in the stroked mice, and (ii) treatment with LM11A-31, and not vehicle, attenuated this p-tau burden within the ipsilateral hemisphere of stroked mice (Figure 9A, *bottom right graph*). A representative image from sections of APP mutant mice served as a p-tau/AT8 staining positive control (Figure 9A, *bottom left image*).

**Figure 9.**
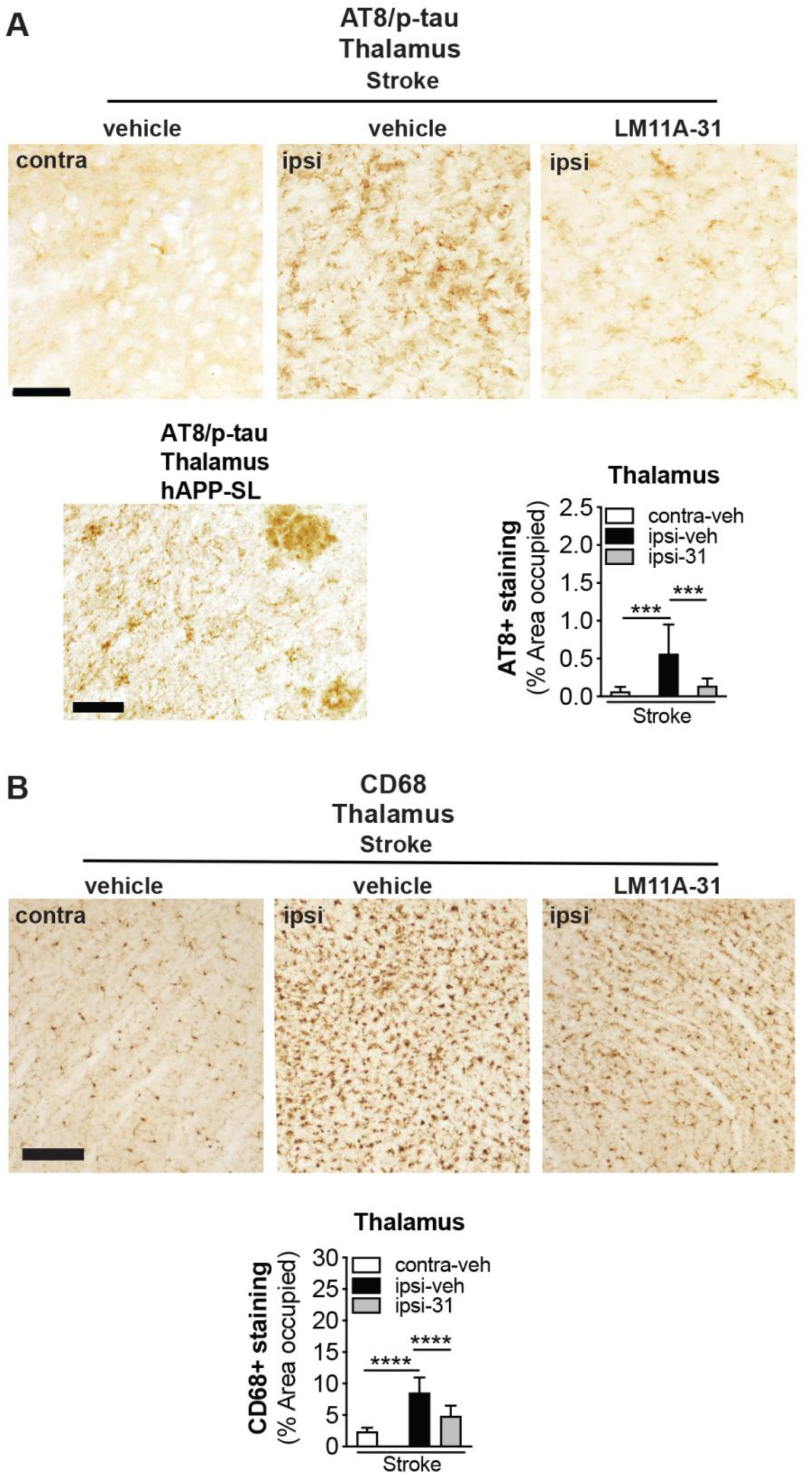
LM11A-31 reduces tau pathology and microglial activation in white matter tracts of the thalamus after stroke. (**A**) Representative images of p-tau labelling using anti-tau phosphorylation (p; AT8) in the thalamus (white matter tracts) of stroked mice treated with vehicle and LM11A-31 (*top images*). Scale bar, 125 μm. A representative image of p-tau labeling in the thalamus of aged hAPP-SL mice is provided as an AT8 staining positive control (*bottom right image*). Scale bar, 50 μm. Quantification revealed significantly more deposits of p-tau in stroked mice treated with vehicle compared to those treated with LM11A-31 (*bottom left graph*). (**B**) Representative 10× images of CD68 labeled microglia and macrophage in the thalamus (white matter tracts) in stroked mice treated with vehicle or LM11A-31 (*top images*). Scale bar, 125 μm. Quantification of CD68 immunoreactivity revealed significantly less staining in LM11A-31 treated stroked mice compared to vehicle treated stroked mice (*bottom graph*). Group sizes: n=10 stroke-veh, n=10 stroke-31. Data represent mean ± SD. ***p<0.001, and ****p<0.0001.

There is evidence for microglial activation in the proximity of tau deposits, and for the interplay of tau pathology and inflammatory cascades (Perea et al., 2020). Stroked mice exhibited numerous CD68+ immunoreactive cells, which displayed an activated microglia morphology (large cell bodies and bushy, thick processes) in the ipsilateral white matter tracks, and a more typical resting morphology (small cell body with long branching processes) in the contralateral white matter tracks. In mice that underwent stroke, LM11A-31 administration decreased the burden of activated microglia (Figure 9B, *top images*). Specifically, quantification of the area occupied by CD68+ labeled microglia/macrophages revealed significantly more CD68+ staining in the white matter tracks of the ipsilateral compared to the contralateral hemisphere of the stroked mice (Figure 9B, *bottom graph*). However, following stroke, LM11A-31 treated mice had less CD68+ staining than vehicle treated mice (Figure 9B, *bottom graph*). These findings demonstrate that LM11A-31 reduces microglial activation in thalamic white matter tracts in aged mice after stroke.

### Phase II: Evaluation of LM11A-31 on recovery following stroke

Given the protective effects of LM11A-31 against brain atrophy, neurodegeneration, and loss of critical neurotransmitters, LM11A-31 was evaluated for its ability to improve recovery after stroke. Motor recovery was assessed by gait using the horizontal ladder rung test (Doyle et al., 2012). Previously, following DH stroke, C57BL/6 mice at 5 months of age displayed a motor impairment of the front limb contralateral to the stroke starting at day 1, which continues until 2 weeks post-surgery (Doyle et al., 2012).

As expected, 2 days following stroke, performance on the ladder rung test was impaired, which localized to the contralateral (left) front limb compared to naïve mice at baseline or pre-surgery (Figure 10A, *left images*: screenshots of video recordings to demonstrate a correct foot placement, and two different types of foot fault: an inability to grip the ladder rung with the front paw, and a foot slip). Scores from day 2 were used to stratify the mice into the LM11A-31 and vehicle treatment groups (Figure 10A, *middle graphs:* stratification method). At 2 weeks, 6 weeks, and 10 weeks following stroke, LM11A-31 treated mice performed better than vehicle treated mice on the ladder rung test, as measured by percent correct foot placement (Figure 10A, *left graphs*).

**Figure 10.**
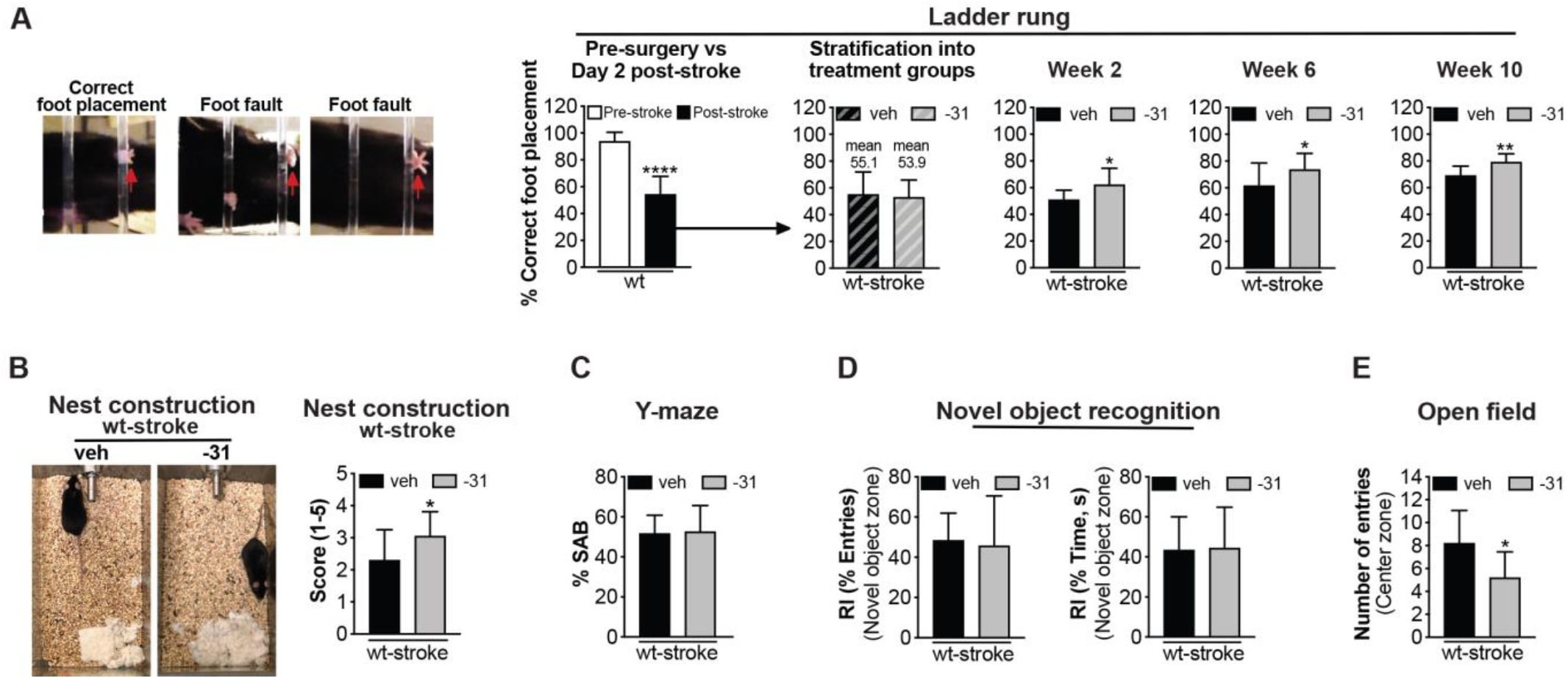
LM11A-31 ameliorates stroke induced impairment of motor function, decline in overall well-being, and anxiety like behavior in aged mice. (**A**) Motor ability on the ladder rung test (*left images show examples of correct foot placement and foot faults*) was assessed at 1 week pre-surgery (*left→ first graph*), and at 2 days (*first graph*), 2 weeks (*third graph*), 6 weeks (*fourth graph*), and 10 weeks (*fifth graph*) post-surgery. On Day 2, there was a motor deficit in stroke operated mice. Ladder rung test scores at 2 days post-surgery were used to stratify the stroked mice into treatment groups (*second graph*). Compared to vehicle treated mice, stroked mice given LM11A-31 displayed a significant improvement of motor function. (**B**) Mouse well-being was assessed with the nest construction test. Among stroked mice, LM11A-31 treated mice displayed significantly better nest building ability than mice given vehicle (*left images and right graph*). (**C**) In stroked mice, LM11A-31 treatment resulted in no significant differences in spatial working memory on the Y-maze. (**D**) LM11A-31 treatment in stroked mice also resulted in no significant differences in object recognition deficits, as assessed by the number of entries into the novel object zone (*left graph*) and the time spent in the novel object zone (*right graph*). (**E**) As assessed through an open field test, impulsive like behavior was significantly lower in LM11A-31 treated stroked mice than in vehicle treated stroked mice. Group sizes: n=10 stroke-veh, n=10 stroke-31. Data represent mean ± SD. *p<0.05, **p<0.01, ****p<0.0001.

Building nests is an innate and spontaneous mouse behavior and nest construction can be assessed and used as a test for psychiatric disturbances that suggest a lack of motivation, anxiety and obsessive compulsive behaviors, or pain (Jirkof, 2014). Typically, mice will make nests achieving scores between 3.5 and 4.5, leaving 0.5 g of nestlets unshredded. Mice with hippocampal lesions achieve a median score of 2.0 (Deacon, 2006), which is in line with the score of our mice at 12 weeks following stroke (Figure 10B, *left images*). However, compared to treatment with vehicle alone, treatment with LM11A-31 significantly improved the performance of mice on the nest construction test following stroke (Figure 10B, *right graph*).

The Y-maze was then used to test spontaneous alternation behavior (SAB). This test is based on the innate interest of rodents to explore a new environment. SAB has been shown to be impaired in AD, and Knowles and colleagues have demonstrated that LM11A-31 attenuates behavioral deficits on a Y-maze (Faizi et al., 2012; Knowles et al., 2013). A naïve or “normal” C57BL/6 mouse would typically score approximately 65-70% SAB (Faizi et al., 2012; Knowles et al., 2013). However, following stroke, we did not detect a difference in SAB between stroked mice given LM11A-31 and those given vehicle (Figure 10C).

Novel object recognition (NOR) is a form of explicit memory known to be impaired in AD. The NOR test was used to evaluate whether LM11A-31 affects AD associated cognitive deficits in the stroke model. Following stroke there was no difference between stroked mice given LM11A-31 or vehicle; both experimental groups spent the same amount of time exploring a novel object relative to a familiar object (Figure 10D).

The open field (OF) test provides a measure of anxiety and impulsivity. In a similar test of the anxiety-impulsivity spectrum, the light-dark transition test, aged, stroked mice display more impulsive risk-taking behavior (Nguyen et al., 2018). Here, we found a difference between stroked mice treated with LM11A-31 and those administered vehicle. Compared to vehicle treated mice, LM11A-31 treated mice entered the open center zone significantly less than the open zone around the perimeter. The vehicle treated mice displayed less cautious behavior, entering the center zone with greater frequency (Figure 10E). This provides evidence that treatment with LM11A-31 mitigates impulsive behavior in stroked mice. Together these data demonstrate that LM11A-31 can promote motor recovery, restore overall well-being, and mitigate impulsive like behavior in aged mice following stroke.

## DISCUSSION

In this study, we asked whether LM11A-31, a small molecule modulator of p75^NTR^ signaling, could be used as a treatment for inhibiting neuronal degeneration, and thereby preserve neuronal health and function after stroke. Our rationale was that decreases in the ratio of neurotrophic versus neurodegenerative signaling play a critical role in other neurodegenerative diseases, such as AD, and studies using mouse models of AD have found therapeutic benefits from treatments targeting p75^NTR^ (Yang et al., 2008; Nguyen et al., 2014). LM11A-31 and its modulation of p75^NTR^ signaling targets ubiquitous pathways and molecules shared among stroke and AD; for example, it can inhibit degenerative signaling via the tau kinases GSK3β, calpain, cdk5, c-jun, and p38, and promote survival signaling via AKT and CREB.

Furthermore, there is evidence that pharmacological modulation of Trk, p75^NTR^, and NGF may prevent memory impairment in a rat model of global ischemia (Choucry et al., 2019), and that neurotrophins, such as BDNF may have therapeutic potential in protection against neurodegeneration and recovery from stroke and traumatic brain injury (Houlton et al., 2019). Following stroke, the expression of both pro-apoptotic and anti-apoptotic signaling molecules and transcription factors that are modulated by LM11A-31 are also upregulated in the ischemic penumbra (Yang et al., 2008; Nguyen et al., 2014; Simmons et al., 2016; Uzdensky, 2019; Yang et al., 2020a).

In the first phase of our study, we found that aged C57BL/6 male mice exhibit changes in brain metabolism following stroke that were attenuated after 2 weeks of oral administration of LM11A-31. To ensure we were evaluating the impact of LM11A-31 on stroke recovery rather than on infarct size, treatment was delayed until 1 week after stroke. One of the most striking changes was in the levels of several neurotransmitters. Stroke induced significant decreases in levels of the neurotransmitter’s glutamate, serotonin, and acetylcholine, and a trend towards a reduction in dopamine. However, the decreases in serotonin and acetylcholine were not seen in the LM11A-31 treated stroked mice, which had significantly higher levels of these neurotransmitters than stroked mice treated with vehicle alone. LM11A-31 treated stroked mice also had significantly higher levels of dopamine than stroked mice treated with vehicle alone.

Serotonin’s biological function is pleiotropic, as it modulates mood, cognition, reward, and learning and memory (Bacque-Cazenave et al., 2020). Acetylcholine is equally important, playing a role in attention, motivation, arousal, and memory (Sofuoglu and Mooney, 2009). The stroke induced reduction in levels of these essential neurotransmitters may impede proper signaling and thereby affect cognition, well-being, motor function, and psychological health. The preservation of the levels of these neurotransmitters and the increase in the levels of dopamine in response to LM11A-31 treatment suggests a broad improvement in neuronal health and function after stroke.

In further support of this improvement, treatment with LM11A-31 for 12 weeks, with administration again delayed until 1 week after stroke, resulted in the preservation of brain volume and prevented loss of NeuN+ immunoreactivity. Atrophy of the brain likely results from progressive degeneration of neurons, and the loss of these neurons likely impedes recovery and contributes to the manifestation of dementia after stroke in some patients.

LM11A-31 may preserve and optimize the function of some neurons better than others, as suggested by our finding that LM11A-31 reduced the loss of ChAT+ immunostaining, but not TH+ immunostaining following stroke. This differential preservation may be explained by varying densities of p75^NTR^ within the cell membranes of different neuronal subtypes; cholinergic neurons are rich in p75^NTR^ and thus may bind more LM11A-31. Attesting to the protective effect of LM11A-31 on cholinergic neurons, other studies have found that treatment with LM11A-31 attenuates cholinergic degeneration in AD mouse models (Knowles et al., 2013; Nguyen et al., 2014; Simmons et al., 2014), and more recently that it suppresses age related basal forebrain cholinergic neuronal degeneration (Xie et al., 2019).

Importantly, in conjunction with its effect on neuropathology following stroke, LM11A-31 also significantly attenuated impairments in motor function on the ladder rung test and significantly improved an activity of daily living in mice, as assessed by the nest construction test. Although the motor deficit induced by the stroke may have affected the ability of the mice to form nests, the improvement in nest building behavior nevertheless demonstrates that LM11A-31 treatment enhanced the ability of the mice to perform an innate behavior associated with their quality of living. LM11A-31 also significantly lessened stroke induced impulsive behavior in the open field test. Notably, in the open field test the vehicle treated mice entered the center zone of the open field arena with greater frequency than the LM11A-31 treated mice. This suggests that their motor disability was not a confound for this test. Together, our data show that the administration of a p75^NTR^ ligand increases neurotransmitter levels, reduces chronic stroke neuropathology, and improves recovery after stroke.

With regard to the mechanisms by which LM11A-31 improved stroke outcome, although LM11A-31 did not affect inflammation in the infarct, it did attenuate deficits in redox homeostasis following stroke by increasing the pool of reduced glutathione available to combat oxidative stress. Furthermore, treatment with LM11A-31 resulted in higher levels of acetyl CoA following stroke, suggesting LM11A-31 mitigated a stroke induced reduction in glycolysis. Although our data do not reveal the mechanism by which acetyl CoA levels were increased by LM11A-31 treatment, an increase in several key metabolites in the glycogenesis pathway suggest that it may be via glycogenolysis. However, further investigation into chronic changes in energy metabolism after stroke and how LM11A-31 treatment affects these changes is required.

We also determined that LM11A-31 treatment attenuated the accumulation of hyperphosphorylated tau in degenerating thalamic white matter tracts of aged mice following stroke. This finding is consistent with the demonstrated ability of LM11A-31 to inhibit the tau kinases glycogen synthase kinase 3 beta (GSK3β), calpain, cyclin dependent kinase 5 (cdk5), c-jun, and mitogen-activated protein kinase p38 (Yang et al., 2008; Nguyen et al., 2014; Yang et al., 2020b). Moreover, there is evidence for microglial activation in the proximity of tau deposits, and for an interplay between tau pathology and inflammatory cascades (Perea et al., 2020). Previous studies have found that LM11A-31 can attenuate increased microglial activation in APP mutant mice with Aβ_42_ and p-tau deposition (Nguyen et al., 2014). In line with this evidence, we observed that LM11A-31 administration decreased microglial activation in the thalamic white matter tracts of aged mice following stroke.

This study was designed with particular attention to STAIR recommendations (Stroke Therapy Academic Industry, 1999; Lapchak et al., 2013; Lapchak et al., 2019; Savitz et al., 2019). Experiments were performed and analyzed in a randomized and blinded manner, we calculated the sample size based on a power analysis prior to the start of the study, and we considered the best route of administration for LM11A-31; oral delivery. We also chose a clinically useful therapeutic window by delaying treatment by 1 week, and selected an appropriate stroke model for evaluating post-stroke recovery rather than acute neuroprotection. However, there are further STAIR recommendations that should be followed prior to implementation of clinical testing. For example, follow up studies are necessary to determine if LM11A-31 can also improve recovery in a stroke model where there is recanalization and reperfusion to emulate patients treated with TPA and mechanical thrombectomy. LM11A-31 should also be tested by a second laboratory, in an alternative mouse strain (e.g., BALB/c mice), in a second species (e.g., rats), and in models that also feature common stroke comorbidities. It also needs to be ascertained if LM11A-31 is similarly effective in female mice. However, with regard to sex, LM11A-31 is similarly effective at reversing pathology in males and females in a mouse model of AD (Simmons et al., 2014), protects against neurodegeneration in both male and female HIV gp120 transgenic mice (Xie et al., 2021), and also protects against cisplatin induced peripheral neuropathy in both male and female mice (Friesland et al., 2014). Therefore, a sex difference is not expected.

A limitation of this study is that the precise mechanism by which LM11A-31 reduced brain atrophy, preserved NeuN+ and ChAT+ immunoreactivity, and improved recovery following stroke is unclear. In AD, LM11A-31 prevents neuronal death, promotes neurotrophin signaling, and inhibits neurodegenerative signaling through the p75^NTR^. Although, there are distinct differences between stroke and AD, the loss of neurotrophic support in the brain is a fundamental neurodegenerative mechanism common to both diseases. However, follow up studies using p75^NTR^ conditional knockout mice are necessary to determine the role of LM11A-31/p75^NTR^ signaling in cholinergic and other specific neuronal sub-populations following stroke.

In conclusion, the findings of this study support a promising role for p75^NTR^ as an effective target in the treatment of stroke beyond the acute setting, for which our current treatment options are extremely limited. Furthermore, because LM11A-31 has already reached Phase IIa of clinical testing as a therapy for AD, once the STAIR recommendations for preclinical testing have been fulfilled, it could rapidly move to clinical trials in patients suffering from stroke and from patients suffering from AD and stroke.

## Acknowledgements

We are grateful to Metabolon for their help with design, data acquisition and data analysis of the metabolomic experiment.

## Authorship contributions

Participated in research design: TVN, FML, KPD

Conducted experiments: TVN, RHC, KEC, FGG, JCZ, MATG, DAB, HWM, SG, JBF

Contributed new reagents or tools: TY, FML

Performed data analysis: TVN, RHC, KEC, HWM, KPD

Contributed to the writing of the manuscript: TVN, RGS, FML, KPD

## List of nonstandard abbreviations

Aβ: Amyloid beta
AD: Alzheimer’s Disease
AKT: Ak strain transforming (RAC alpha serine/threonine-protein kinase)
APP: Amyloid precursor protein
BDNF: Brain derived neurotrophic factor
Cdk5: Cyclin dependent kinase 5
ChAT: Choline acetyl transferase
DH Stroke: Distal middle cerebral artery occlusion + hypoxia stroke
dMCAO: distal Middle cerebral artery occlusion
FO: Familiar object
G-CSF: Granulocyte-colony stimulating factor
GM-CSF: Granulocyte macrophage-colony stimulating factor
GSH: Reduced glutathione
GSK3β: Glycogen synthase kinase 3 beta
GSSG: Oxidized glutathione
IFN-γ: Interferon gamma
IL: Interleukin
IP-10: Interferon induced protein-10
IRAK: Interleukin-1 receptor associated kinase
KC: Keratinocytes derived chemokine
MCP-1: Monocyte chemoattractant protein-1
MIP-1α: Macrophage inflammatory protein-1 alpha
MIP-1β: Macrophage inflammatory protein-1 beta
MIP-2α: Macrophage inflammatory protein-2 alpha
NeuN: Neuronal nuclei
NFκB: Nuclear factor kappa-light-chain-enhancer of activated B cells
NGF: Nerve growth factor
NMDA: N-methyl-D-aspartate
NO: Novel object
NOR: Novel object recognition
OF: Open field
PCA: Principal component analysis
P75^NTR^: p75 neurotrophin receptor
PI3K: Phosphoinositide 3 kinase
RANTES: Regulated on activation normal T cell expressed and secreted
RI: Recognition index
SAB: Spontaneous alternation behavior
TCA: Tricarboxylic acid
TH: Tyrosine hydroxylase
Thy1: Thymocyte differentiation antigen 1
TNF-α: Tumor necrosis factor-alpha
TPA: Tissue plasminogen activator
UPLC-MS/MS: Ultrahigh performance liquid chromatography/tandem mass spectrometry

## Recommended section assignment

Drug Discovery and Translational Medicine

## FOOTNOTES

This work was supported by the National Institutes of Health National Institute of Neurological Disorders and Stroke [Grant R01NS096091], the National Institute on Aging [Grants R01AG063808 and R21AG062781], the United States Department of Veterans Affairs [Grant I01RX003224], the Jean Perkins Foundation and by the Leducq Foundation [Grant Stroke-IMPaCT: Stroke – immune mediated pathways and cognitive trajectory].

